# Mutational profile of the regenerative process and *de novo* genome assembly of the planarian *Schmidtea polychroa*

**DOI:** 10.1101/2023.07.20.549885

**Authors:** Ádám Póti, Dávid Szüts, Jelena Vermezovic

## Abstract

Planarians are organisms with a unique capacity to regenerate any part of their body. New tissues are generated in a process that requires many swift cell divisions. How costly is this process to an animal in terms of mutational load remains unknown. Using whole genome sequencing, we defined the mutational profile of the process of regeneration in the planarian species *Schmidtea polychroa*. We assembled *de novo* the genome of *S. polychroa* and analyzed mutations in animals that have undergone regeneration. We observed a threefold increase in the number of mutations and an altered mutational spectrum. High allele frequencies of subclonal mutations in regenerated animals suggested that relatively few stem cells with high expansion potential regenerated the animal. We provide, for the first time, the draft genome assembly of *S. polychroa*, an estimation of the germline mutation rate for a planarian species and the mutational spectrum of the regeneration process of a living organism.

## INTRODUCTION

Living organisms differ significantly in their capacity to regenerate damaged or lost tissues. Some organisms, like planarians, are capable of rebuilding a complete body from a small number of cells, while others, like *C. elegans* cannot replace damaged tissues at all. In mammals, regeneration is restricted, with the liver being particularly efficient in eliciting tissue repair response, but also other tissues have a limited potential to regenerate [1]. The process of regeneration requires a complex orchestration of basic biological processes: proliferation, differentiation, and cell death, in a very similar way as during embryogenesis or in pathological conditions like carcinogenesis. A physiological process like embryogenesis is not cost free at the level of the genome of proliferating cells. Mutation rates of fetal tissues can exceed fivefold the mutation rates of the same tissues over a year period in an adult [2]. In fact, mutations acquired during embryogenesis are a risk factor for development of cancer later in life [3].

Easier accessibility to whole genome sequencing has made it possible to study mutational spectra of physiological and pathological processes in great detail over the last decade. Human *de novo* germline mutation rates and their spectra have been determined [4], as well as somatic mutation spectra of specific tissues across different mammalian species [5] or somatic mutations in collections of cancer tissues [6]. All these processes leave mutational “scars” at the genome level, with smaller or greater consequences to the functioning of an organism. In a lifetime of an individual, tissues will be exposed multiple times to mechanical, chemical or in some cases surgical insults, that elicit tissue repair response, but the potential mutational burden of this process has not been assessed yet. To answer this question, we chose the most efficient and accessible model system for regeneration studies – the planarian. Regeneration in planarians is an extremely fast and accurate process which can produce a new individual from a small piece of a tissue, amputated from any body part. Abundance of pluripotent somatic stem cells (traditionally referred to as neoblasts) and a complex system of positional information underlie this process, which gained a solid mechanistic framework in the last two decades [7–9]. It takes only about seven days for a fragment of an animal to regenerate missing tissues, increasing numbers of proliferating stem-cells fivefold within 48 hours [10].

We have previously published a pipeline based on an algorithm – IsoMut – for the analysis of unique mutations in isogenic whole-genome datasets [11]. IsoMut was used successfully to elucidate the mutational signature of cisplatin, and the mutational profiles of several other chemotherapeutics or genetic treatments, in the context of oncological studies [12–15]. Here, for the first time, we applied it to a whole organism – the planarian *Schmidtea polychroa* (*S. polychroa*), to study the mutation rate of a physiological process, regeneration. *S. polychroa* is surprisingly easy to maintain and propagate in the lab, but its main advantage for the scope of this study was the ability to reproduce by parthenogenesis – which satisfies the requirement for the isogenic background. It has been observed that triploid biotypes of *S. polychora* originating from Western Europe reproduce by pseudogamous parthenogenesis [16].

*S. polychroa* is not a new model in regeneration studies. Several versatile and interesting biological questions have been addressed in this system. Mechanisms of planarian embryogenesis, unique among metazoans and characterized by a divergent gastrulation stage with temporary feeding structures, were studied in *S. polychroa* [17]. Another notable aspect of the physiology of *S. polychroa* is the absence of aging, as measured by mass-specific metabolic rates [18]. Evolutionary studies have addressed the intricate question of successful survival of a parthenogenetic species, which is rescued by rare, occasional sexual reproduction [19].

Though *S. polychroa* shows a number of advantages as a model system, the main drawback remains the lack of a published reference genome. To bridge this gap and to answer whether regeneration is associated with a genome-wide increase in DNA mutations, we set out to sequence and assemble the genome of *S. polychroa*. Using protocols for isolation of high-quality planarian genomic DNA, PacBio long-read sequencing, followed by *de novo* genome assembly, we provide the first draft genome of *S. polychroa*. We describe the structure of the genome and validate it through comparison with the well characterized species *Schmidtea mediterranea* (*S. mediterranea*). Finally, whole genome sequencing of parent-regenerant-parthenogenetic offspring trios allowed us to estimate the mutation rate of *S. polychroa*, calculate the mutational burden and spectrum associated with the regeneration process, and show that stem cells contribute differentially to regeneration.

## RESULTS

### *De novo* assembly of a novel planarian genome

Adult *S. polychroa* animals range 1–2 cm in size. Their epidermis is of dark brown pigmentation and adult animals develop multiple photoreceptors (Fig 1A). *S. polychroa* is a cross-fertilizing, pseudogamous hermaphrodite, meaning that the sperm of one animal activates embryogenesis of the egg of another animal, but the genetic material of the first animal is excluded from the zygote [20]. Oocytes of these animals pass through a premeiotic doubling of the triploid genome, followed by an asynaptic meiosis without effective recombination of the genetic material [21]. Though *S. polychroa* reproduces mainly by parthenogenesis, occasional sex has been observed in this species at very low frequency [19].

**Figure 1.**
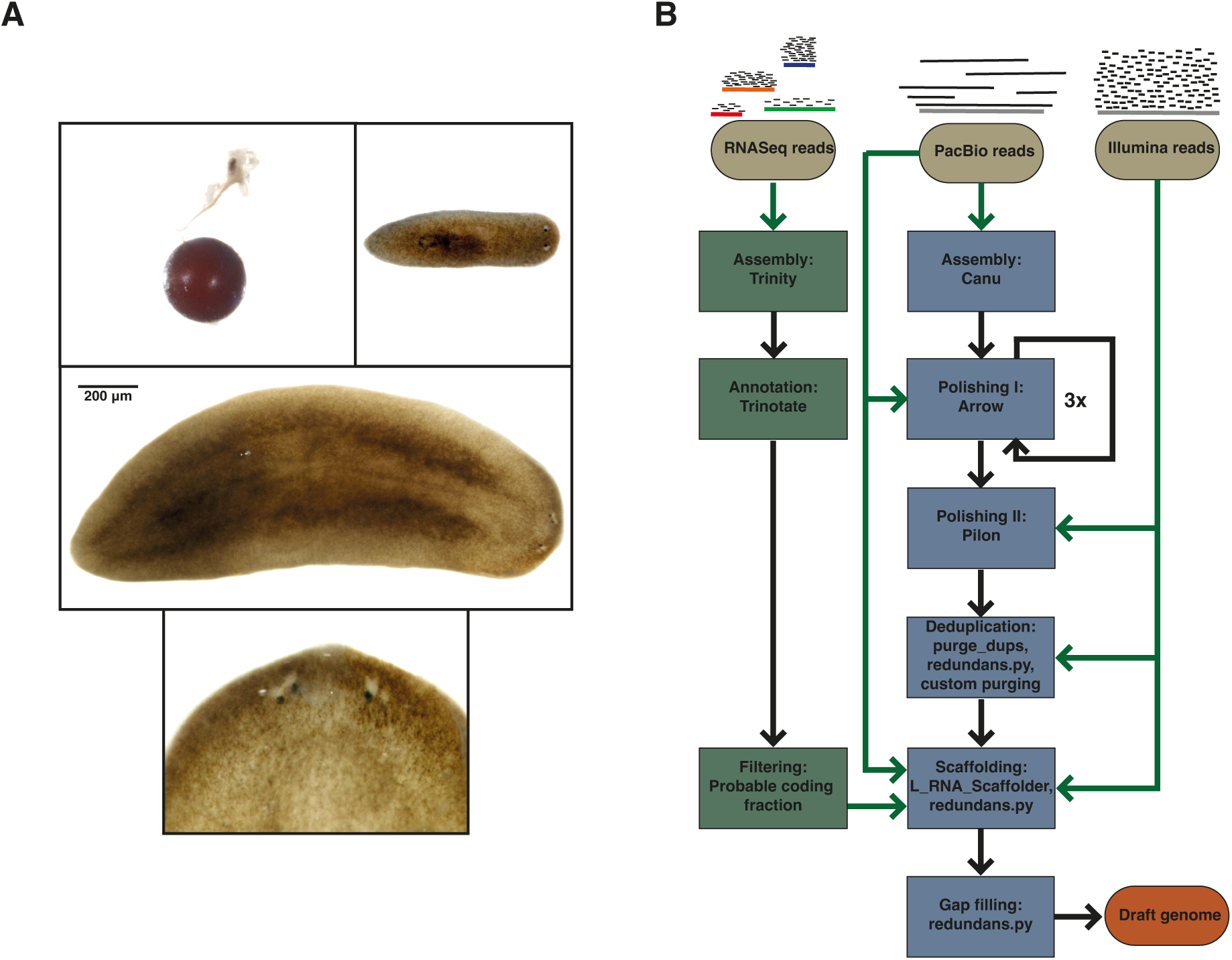
*De novo* genome assembly strategy. **A)** Images of *S. polychroa*; cocoon (top right), young (top left) and adult animal (middle), close up on the head region of the adult indicating multiple photoreceptors (bottom). **B)** Flowchart of the *de novo* genome assembly strategy.

To obtain high-quality genomic DNA, required for long-read PacBio sequencing, we isolated genomic DNA from 20 animals according to a previously published protocol for the extraction of genomic DNA from *S. mediterranea* [22]. For the assembly of the draft genome, we generated 18.04 Gbp of PacBio long reads with an average read length of 15.24 kbp and 39.42 Gbp of 150 bp paired-end Illumina reads. As planarian genomes are generally GC-poor, we assembled the long reads with Canu using parameters optimized for lower coverages and skewed GC-ratios. After polishing with Illumina and PacBio reads, scaffolding and haplotig purging (Fig 1B, Fig S1), our draft genome, referred below as bm_Spol_g1, consisted of 8253 scaffolds with an overall genome length of 434.3 Mbp and an N50 value of 80.7 kbp.

Wild populations of *S. polychroa* represent at least four coexisting biotypes [20] with four chromosomes, different ploidy levels and experimentally determined haploid genome sizes in the range of 0.56-1.29 Gbp [23]. To characterize the biotype of our animals, we estimated the true genome size using several approaches (Fig 2A, Table S1): k-mer based estimations with Genomescope2, a coverage-based measure using an aligned Illumina dataset and the Lander-Waterman equation [24], and gene content-based approaches according to BUSCO [25]. The average haploid genome size was 587.9 Mbp, indicating that our draft assembly was ∼74% complete. In addition, we measured genome size experimentally by flow cytometry [26], comparing the fluorescent signal of propidium-iodide stained nuclei of *S. polychroa* to that of a species of a known genome size, *S. mediterranea*. 588.67 Mbp obtained by flow cytometry correlated very well with the theoretical value of 587.9 Mbp (Fig 2B, Fig S2, Table S2). To confirm the ploidy level of the biotype used in the study, and to exclude the possibility of paternal contribution to the genetic material through occasional sex, we performed an analysis of allele frequency distribution of high-confidence germline mutations. It has been reported that presence of tetraploid offspring can be a good estimator of occasional sex rate in *S. polychroa* [27]. The distribution of allele frequencies of germline mutations showed peaks at 33% and 66% (Fig 2C), indicative of a triploid genome, which was also supported by the presence of 3 peaks on the KAT profile of the reference genome (Fig S2B). Triploidy was similarly confirmed for all animals used in this study (Fig S3).

**Figure 2.**
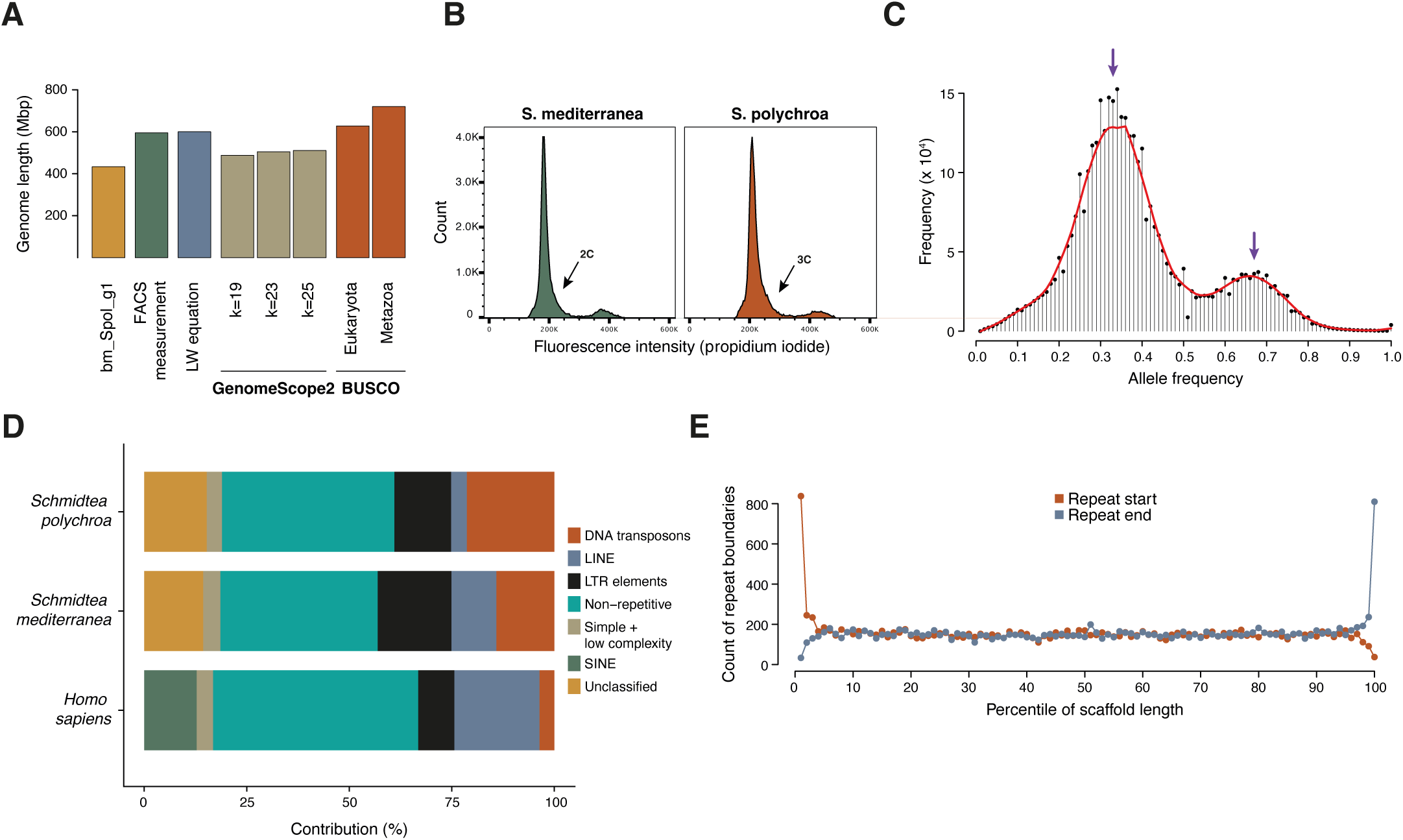
Analysis of genomic features in the *S. polychroa* assembly. **A)** Estimation of genome size by multiple methods. Each bar represents a genome size estimation according to the indicated approach. **B)** Representative FACS profiles of *S. polychroa* and *mediterranea* cells, stained for their DNA content. **C)** Histogram of allele frequencies of germline mutations. The two peaks of the distribution at 33% and 66% refer to the major and minor alleles at a heterozygous site and indicate that the *S. polychroa* strain used in theis study was triploid. **D)** Distribution of repeat element classes in the genomes of *H. sapiens, S. polychroa* and *S. mediterranea.* **E)** Histogram of relative positions of repeat elements. Only DNA elements longer than 2 kb were considered, and scaffolds lengths were normalized to 1. The individual points refer to the number of repeat instances having their start (red) or end (blue) in the respective percentile of the normalized scaffold length.

The draft genome of *S. polychroa* contained 57.98% of repetitive elements as determined by RepeatModeler and RepeatMasker (Table S3). Indeed, planarian genomes are known to have high levels of repetitive DNA content [22]. The distribution of different elements showed a profile differing from vertebrate genomes and had an especially high level of DNA elements and unclassified repeats, similar to the repeat content of *S. mediterranea* [22] (Fig 2D). The high prevalence of repetitive elements contributed to the fragmentation of the draft genome: elements longer than 2 kbp were overrepresented at the ends of scaffolds (Fig 2E).

Next, we assembled the mitochondrial genome of *S. polychroa* (Fig S4). We detected non-canonical start codons, with TTG being used in 6 out of 13 mitochondrial protein coding ORFs. The gene order in the mitochondrial genome of *S. polychroa* is highly conserved with respect to the other members of triclads. The genome is interspersed with long non-coding regions that increase the length of the mitogenome to a surprising 25534 bp. Similar features of mitochondrial genomes have been described in other *Platyhelminthes* [28].

### The genomic architecture of *S. polychroa*

To have an independent measure of the completeness of our genome assembly, we generated a *de novo* transcriptome assembly using 59.8 million Illumina 150PE reads pairs, utilizing RNA isolated and sequenced from multiple *S. polychroa* individuals; we refer to this assembly as bm_Spol_tr1. We evaluated several quality control measures of the bm_Spol_tr1 transcriptome and found that the assembly contains the information of over 98% of the RNA-seq reads (Fig S5A-B), and in terms of BUSCO content and number of genes/transcripts it is comparable to the already existing transcriptome assembly available on PlanMine [29] (Fig S5C-D).

Transcripts were back-mapped onto the bm_Spol_g1 genome assembly with GMAP [30] (Fig 3A). For the back-mapping, we used only the longest isoforms of the probable coding fraction (PCFL set) from the bm_Spol_tr1. Out of 25054 transcripts in the PCFL set, 17735 transcripts were mapping at a single locus, 4960, or 19.79% were not mapped by GMAP, in agreement with our previous estimate of more than 80% completeness of the bm_Spol_g1 assembly, and the remaining transcripts were mapped as duplicates, at multiple positions or in two chimeric fragments; these transcripts were further analysed manually (Table S4). Almost all multi-mapper transcripts overlapped with RepeatModeler models, suggesting that these are mostly transcribed repeat elements. In the case of double mappers, 51.6% of the transcripts overlapped with repeat elements, 30.8% were in fact fragmented, and 7% were presumably real duplications, mostly mapped adjacently on the same scaffolds. Importantly, only 37 out of the 25054 PCFL transcripts were remaining unpurged haplotigs on two different scaffolds, and upon visual inspection these were never longer than a few kbps. In accordance with the analysis of k-mer abundances (Fig S1B), this proves that heterozygous regions were appropriately collapsed. Among the chimeric transcripts, i.e., those mapping to non-overlapping fragments on two different scaffolds, repeat-overlapping transcripts were also present with a high contribution of 41.3%, while the remaining chimeric mappings were caused either by spliced-leader trans-splicing (12.5%), true fragmentation of bm_Spol_g1 (38.3%), erroneous scaffolding (1.8%) or incorrect GMAP mapping of short transcript fragments to nearby homologous regions (6%). Trans-splicing used a spliced-leader sequence (Fig S6), encoded by 6 spliced-leader genes in several clusters as assessed by SLIDR [31], and both the sequence and the secondary structure of this non-coding RNA was similar to spliced-leaders identified in *S. mediterranea* [32].

**Figure 3.**
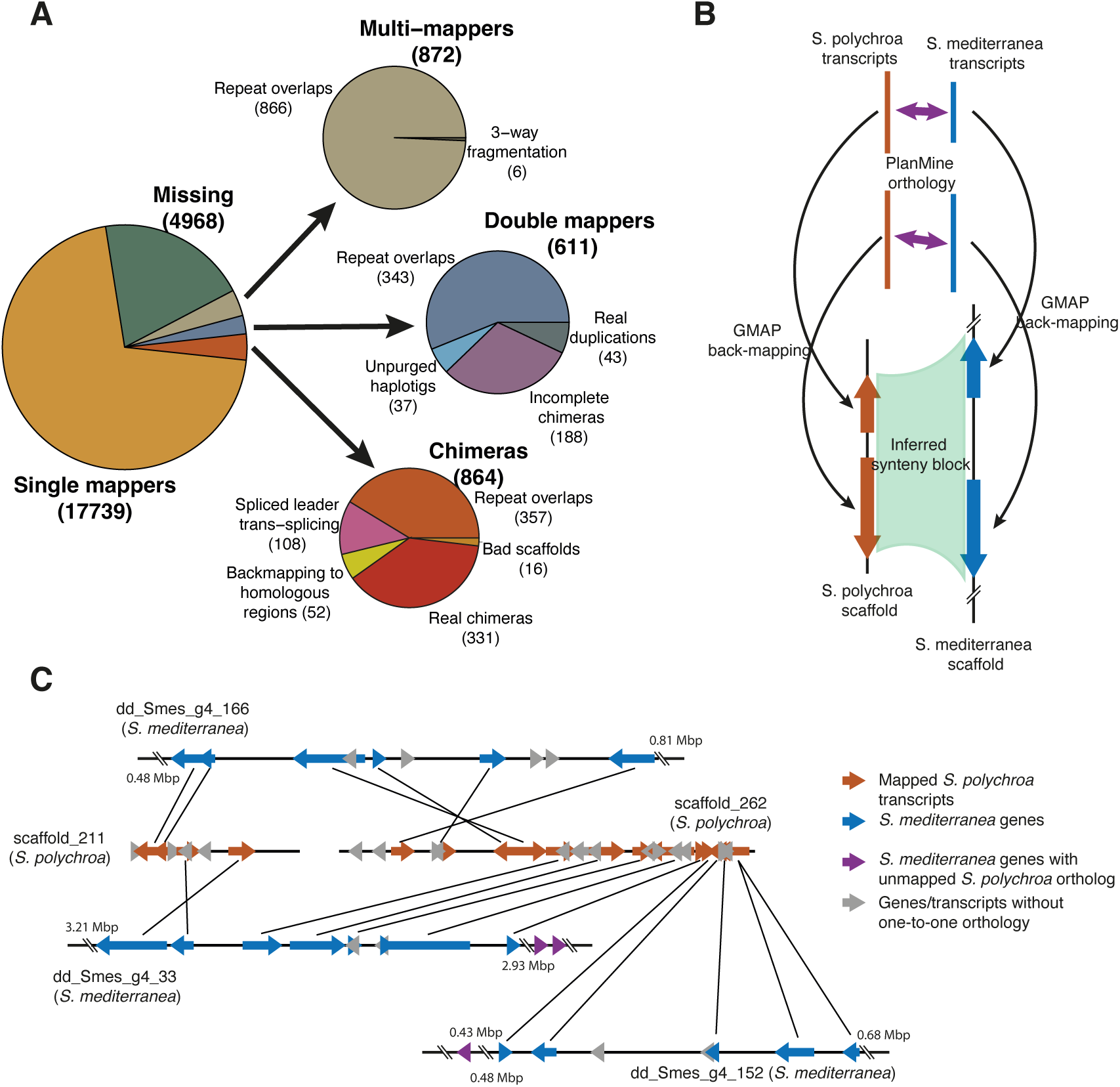
Genome characterization. **A)** Analysis of transcriptome back-mapping. Automatic annotation by GMAP was manually revised to enable finer categorization. The majority of abnormal transcript mappings are due to overlaps with repeat elements. **B)** Scheme of synteny identification. *S. polychroa* and *S. mediterranea* trancripts (orange and blue ribbons, respectively) were mapped to the respective genome assemblies by GMAP (thin black arrows), and interspecies homologies were taken from PlanMine (thick purple arrows). **C)** A representative example of synteny regions between the two *Schmidtea* species. Back-mapped transcripts on *S. polychroa* scaffolds *scaffold_211* and *scaffold_262* (orange ribbons) uncover a reciprocal translocation between to *S. mediterranea* scaffolds *dd_Smes_g4_166* and *dd_Smes_g4_33*. Some mapped genes on scaffold_262 are also linked to a third S. mediterranea scaffold, *dd_Smes_g4_152*: here the exact breakpoint cannot be resolved, because relevant *S. polychroa* transcripts in the region are not mapped on the bm_Spol_g1 assembly (purple ribbons).

Next, we tested the quality of the described genomic architecture of *S. polychroa* through a comparison with a reference genome of *S. mediterranea*. Using GMAP, we mapped the PCF sets from transcriptomes of both species against the bm_Spol_g1 and the dd_Smes_g4 [22] assemblies, respectively, and used homology information from PlanMine [29] between the two *Schmidtea* species to predict synteny regions using only one-to-one homologies (Fig 3B). Altogether, 4434 scaffolds were linked only to one *S. mediterranea* scaffold, suggesting direct synteny between the regions (Fig S7A). These synteny blocks had a mean length of 17624.5 bp, with the maximal length being 427088 bp, indicating that the identification of synteny regions was limited by the remaining fragmentation of the bm_Spol_g1 assembly (Fig S7B).

We also found 515, 84, 11 and 1 *S. polychroa* scaffolds with 2, 3, 4 or 5 linked *S. mediterranea* scaffolds, respectively. These events either mark potential rearrangement sites between the two species, or erroneous scaffolding. To differentiate between these possibilities, we manually evaluated 157 potential break sites on the bm_Spol_g1 scaffolds where at least two genes were on both sides mapping to different scaffolds in *S. mediterranea* (Fig S7C, Table S5). We found that 22 such rearrangement sites were probably valid events representing the divergence of a single *S. polychroa* genomic segment in *S. mediterranea*: in some cases, the exact breakpoints could be found, revealing reciprocal translocations and large-scale inversions (Fig 3C). We also found that 75 represented poor scaffolding, determined by the presence of gaps with zero coverage in both Illumina and PacBio read alignments, or by finding exclusively very short (<10 bp) overlaps. Interestingly, 49 potential break sites mapped to the very ends of two different *S. mediterranea* scaffolds. Comparison of these regions to a more recent chromosome-level assembly of *S. mediterranea* [33] showed that the sequences of these dd_Smes_g4 scaffolds are indeed positioned adjacently (Table S6), giving an example how the comparative genome analysis of two close relative species can contribute to more complete assemblies.

### *De novo* mutation rate in *S. polychroa*

*De novo* mutation rates show similarities across living organisms; however, *de novo* mutation rate of any planarian species has not been established yet. An effort to measure the number of lineage-specific mutations extracted from transcriptomes of two *D. japonica* strains has been made previously [34], but without success in estimating mutation rates or whether mutations accumulated over time. To determine *de novo* mutation rates in *S. polychroa*, we took advantage of the parthenogenetic reproduction strategy of the triploid biotypes of this species, as only the maternal germline will contribute to the genetic content of the offspring. This way, any heterozygous mutation detected in an offspring, but not found in its parent, will represent the mutagenic events that occurred in time, between the parental zygote and the zygote of the offspring (Fig 4A, control group). This scenario in conceptually similar to the role of single cell cloning steps in induced mutation studies [12].

**Figure 4.**
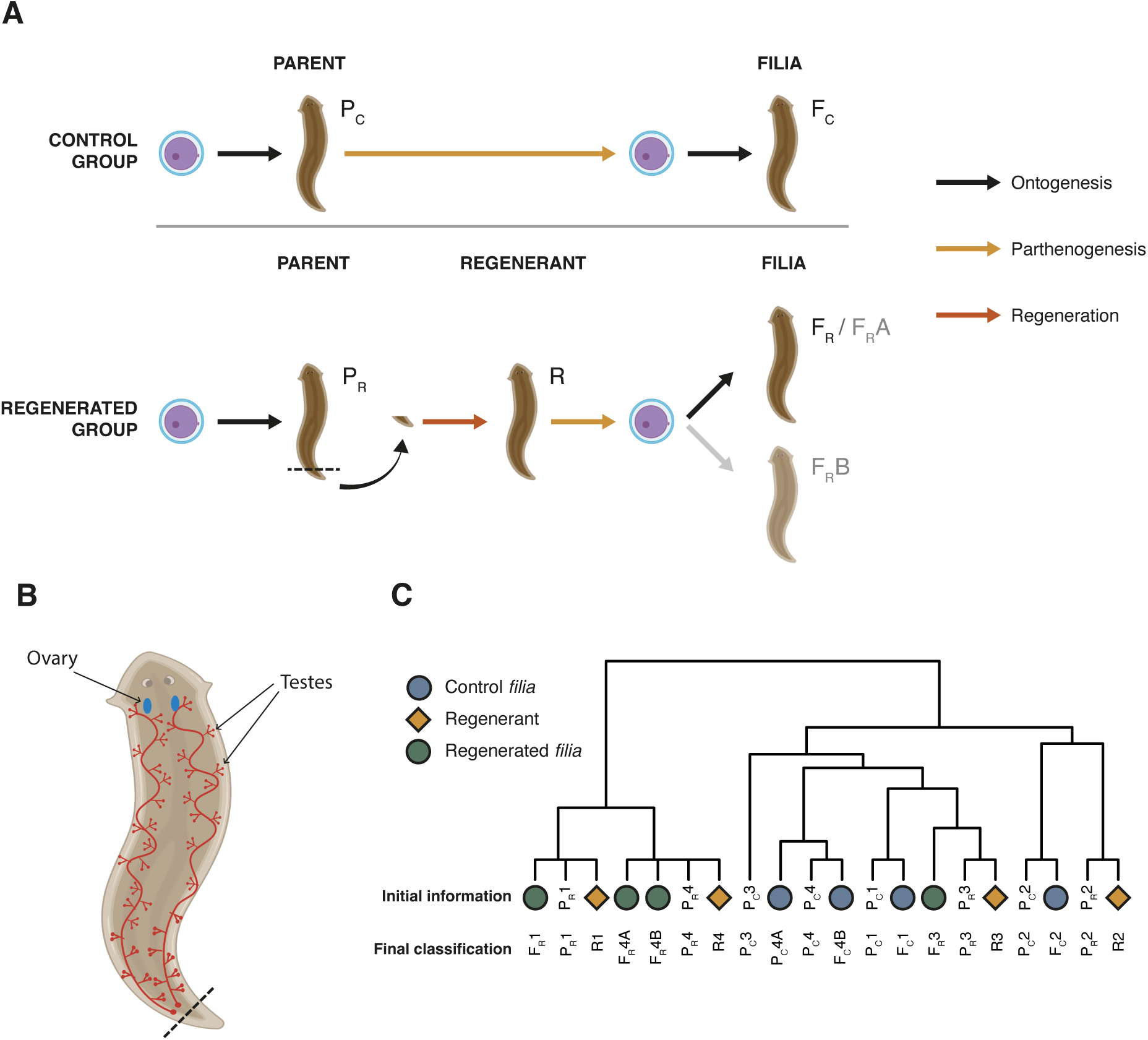
Experimental set up and phylogenetic relations between samples. **A)** Scheme of experimental design for the analysis of *de novo* mutations. For the control group (top half), parental animals (P_C_) were left to reproduce by parthenogenesis (n = 4) and progenies (*filia –* F_C_) were collected and sequenced. In the case of the regenerated group (bottom half), tails of parental animals, or P_R_ (n = 4) were amputated and left to regenerate and reach sexual maturity (R). Regenerants reproduced by parthenogenesis and the progenies (F_R_) were collected for the analysis. The panel was created with bioRender.com. **B)** Schematic figure depicting the positions of the reproductive organs in the planarian body, and the approximate cut site. **C)** Phylogenetic tree showing the planaria lineages used in this study. The colored symbols show the initial information regarding the identities of each sample.

To find *de novo* mutations, we first collected 4 hatchlings from a population of four adult *S. polychroa* (parental generation: P_C_1-P_C_4) and let them to develop into adulthood (Fig 4A). Genomic DNA was isolated from both the parents and the *filia*. After whole genome sequencing, the resulting genomic DNA datasets were aligned against the bm_Spol_g1 draft genome, and point mutations and short indels were called in parallel by IsoMut [11] and HaplotypeCaller [35]. The two methods are different, yet complementary. While IsoMut is a ploidy-agnostic mutation caller that only considers the number of supporting reads in a given sample, and efficiently filters out noise by concurrent analyses of many samples, HaplotypeCaller is a program often used on cancer samples with chaotic ploidy, that works by performing local re-assemblies around potential mutations. Double mutation calling step allowed for more scrupulous detection of real mutations.

As the animals were kept in a shared vessel, it was unknown which offspring belonged to which parent. By analysing shared, lineage-specific mutations between parents and the filial generation, we identified that the four parent animals produced 1, 1, 0 and 2 offspring, respectively (Fig 4C, Table S7). The *filia* were named F_C_1, F_C_2, F_C_4A and F_C_4B, with the numbers referring to the respective parent, and the A/B notation marking siblings from the same parental animal.

The filial generation contained on average 5 +/- 2.94 (mean +/- SD) unique clonal SNVs and 1.25 +/- 1.5 unique clonal indels that were not detectable in the respective parents (Fig 5A, Table S8): these clonal mutations were not found in the parental zygote but were universally present in all cells of the offspring, and thus represent the total mutation load of a generation. By considering only mutations positioned in the uniquely mappable fraction of the assembly, having a length of 303.93 Mbp, and a triploid genome, the *de novo* rates amount to 1.31 x 10^-8^ per base pair per generation for SNVs, or 1.72 x 10^-8^ per base pair per generation overall.

**Figure 5.**
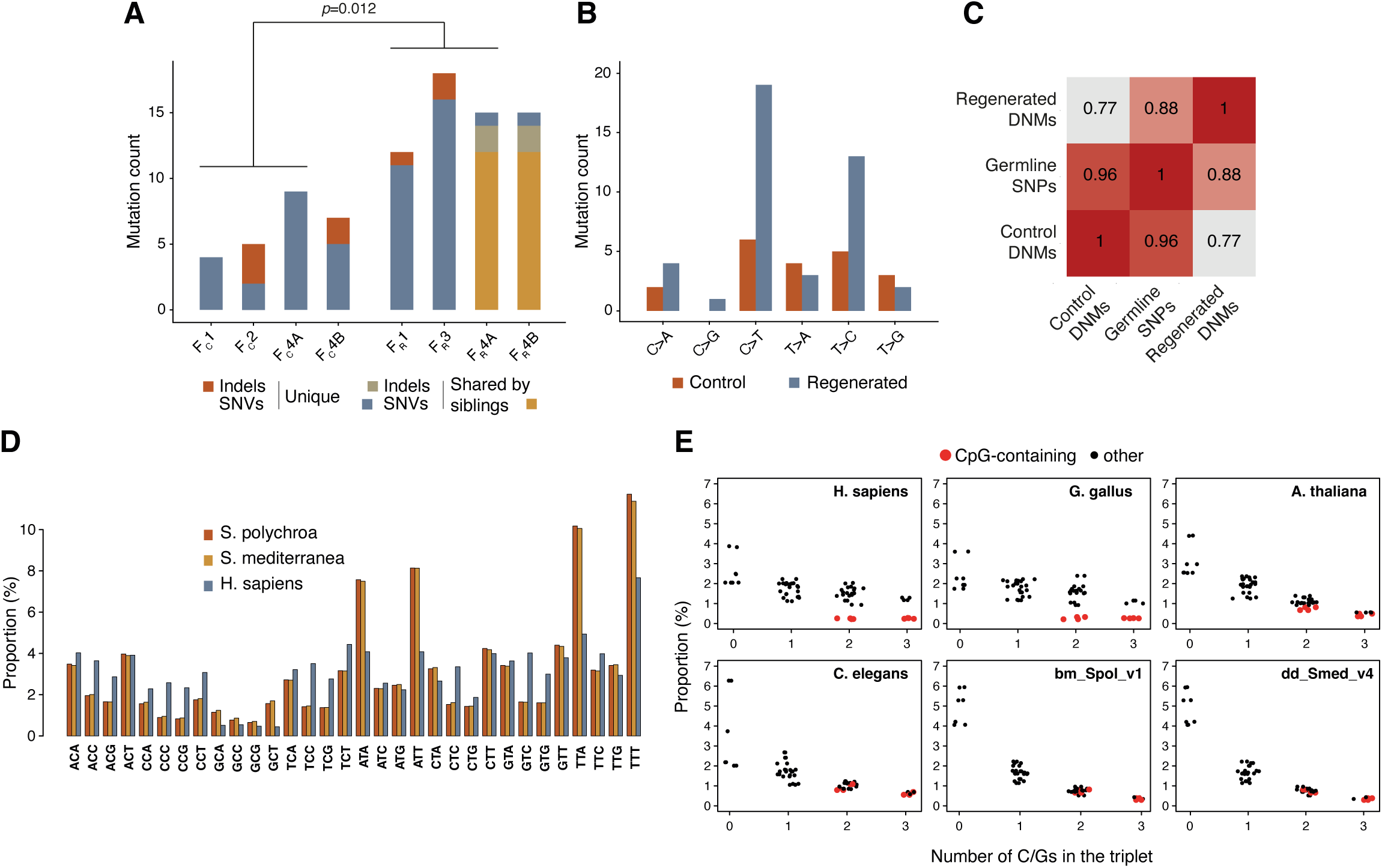
Analysis of *de novo* mutations. **A)** Counts of *de novo* mutations in progeny of regenerated animals versus progeny of control animals. Mutations shared by siblings were marked. Significance was assessed by unpaired two-sample t-test. **B)** Mutational spectra of *de novo* mutations. **C)** Comparisons of Pearson correlations between the spectra of regenerated and control *de novo* substitutions and germline mutations shared by all animals. **D)** Bar plot showing the frequencies of trinucleotides in the genome of *S. polychroa, S. mediterranea* and *H. sapiens.* **E)** Scatter plots indicating the relationship between triplet frequencies and counts of G/C bases in each trinucleotide. Triplets containing CpG motifs are shown in red.

### Regeneration induces additional mutations in filial generation

During regeneration, germ cells arise from the same pool of stem cells as somatic cells [36], suggesting that patterns of mutagenic consequences of whole-body regeneration will also be captured as changes in the genome of the filial generation, in addition to *de novo* mutations accumulated during the parthenogenetic reproductive cycle. We expanded our previous experimental set up (Fig 4A) and started from four different parental animals (P_R_1-P_R_4) that underwent an amputation step prior to parthenogenesis. Most of the animal body was amputated, including ovaries, and only a tip of the tail was left to regenerate into a new individual (Fig 4B). When fragments completed regeneration and developed into adults (R1-R4), they were allowed to lay eggs. Four offspring animals were collected and together with the respective regenerated and parental animals, were used for DNA extraction and whole genome sequencing. Shared mutations in the 12 samples uncovered the relations between the animals (Fig 4C) and assigned a regenerated animal (R) and a parent (P_R_/P_C_) to each of the offspring animals that were analysed. We discovered that while F_R_3/R3 had no descendants among the *filia*, F_R_4/R4 had two offspring, so the filial specimens were named F_R_1, F_R_2, F_R_4A and F_R_4B. When contrasted with the P_R_ parent genomes, we found that the filial generation harboured 13.5 +/- 2.52 unique SNVs and 1.5 +/- 0.58 unique indels. This implied that regeneration caused a nearly threefold increase in the *de novo* SNV mutation rate in one generation (Fig 5A) (*p*=0.012, unpaired two-sided t-test). Interestingly, the majority of mutations were shared in the siblings F_R_4A and F_R_4B. This suggested that these animals developed from germ cells in the R4 regenerant that were descendants of the same stem cell lineage, which probably underwent a high number of cell divisions during the regeneration process prior to germline specification. The same finding also confirmed that the detected clonal filial mutations were present in the respective oocyte and were not generated during embryogenesis.

The point mutation spectra of the *de novo* mutations in control versus regenerated filial generation showed a marked difference between the two groups (Fig 5B). SNVs in control animals had a broad spectrum, while in the regenerated filial generation an elevated number of C>T and T>C transitions could be observed. We compared the *de novo* spectra of control and regenerated animals to the spectrum of lineage-specific germline SNPs, those that were shared by all animals included in this study. We found that the profile of SNPs shared by all animals resembled more closely the *de novo* spectrum of the control group (Fig 5B-C, Fig S8) than that of the regenerated group, implying that regeneration had no contribution to the spectrum of germline mutations in studied *S. polychroa* population, in accordance with previous observations that *S. polychroa* does not reproduce by fission. Confirming this established fact, through comparison of mutational profiles, provides validation for our mutational analysis tool. A general source of C>T mutations in living organisms is methyl-CpG deamination, which is due to the higher propensity of methylated cytosines in the CpG context for spontaneous hydrolysis, resulting in thymine bases at the site [37]. Though DNA methylation has been detected across *Platyhelminthes* [38], no measurable 5-methyl-cytosine levels have been found by the use of methylation sensitive restriction enzymes or antibodies against 5-methyl-cytosine in the planarian *S. mediterranea* [39]. We applied a genome wide approach to test whether cytosines are methylated in *S. polychroa*. The presence of CpG methylation can be deduced by genomic triplet frequencies: CpG containing triplets, being inherently less stable, are underrepresented in highly methylated genomes, like those of vertebrates [40]. To determine the contribution of CpG methylation in the elevated rate of C>T mutations after regeneration, we compared the genome-wide triplet spectra of *S. polychroa*, *S. mediterranea* and *Homo sapiens* (Fig 5D). In accordance with the low GC ratios of planarian genomes, GC-rich triplets are generally depleted, however, CpG-containing triplets were not particularly underrepresented (Fig 5E), as in vertebrate genomes. The profile of planarian CpG-containing triplets was more similar to the non-methylating *Caenorhabditis elegans*, implying that CpG methylation was not present in *S. polychroa*.

### Regenerated animals are mosaics for the regeneration-induced mutations

To understand better the origin of *de novo* mutations after regeneration, we tested whether genomic changes in the filial generation were pre-existing in their ancestors. In the filial generation, altogether half (23/46 in the regenerated group) of the clonal mutations had high-confidence support in the corresponding regenerated animal (Fig 6A, Table S7), with allele frequencies in the range of 1.09 – 5.56%. As this set also included events supported by low number of reads, we validated a subset of these events (32 out of 46) by PCR and high-coverage amplicon sequencing. This approach confirmed the presence of most of these variants (30/39) in regenerated animals, even in cases where the coverage depth of WGS was insufficient to detect them (Fig 6B, Table S8). The allele frequencies detected by the two methods showed a moderate correlation of 0.62. Interestingly, in the case of F_R_4A-F_R_4B sibling pair, shared *de novo* mutations were mostly subclonal in R4 (10/14 by WGS, 71.4%), while the two unique mutations could not be detected in R4 even by the PCR approach, indicating again that the unique mutations, which amounted to approximately 10% of total clonal filial mutations in these two animals, arose later than the shared ones, probably after regeneration was completed.

**Figure 6.**
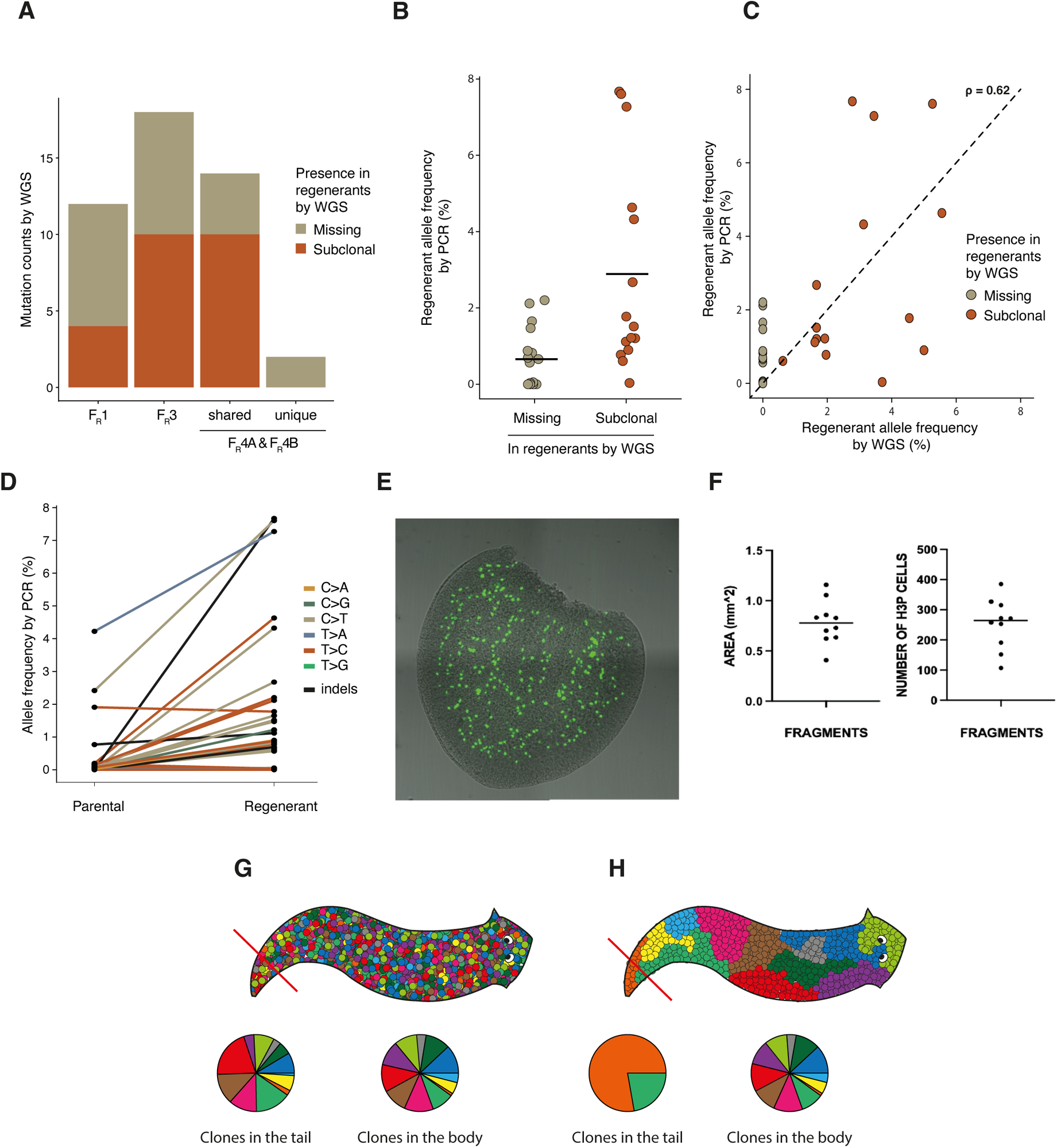
Analysis of pre-existing subclonal mutations in regenerants. **A)** Ratios of mutations found in filia that are also identified as subclonal in the corresponding regenerant. **B)** Allele frequencies of filia clonal mutations detected by amplicon sequencing in the corresponding regenerant animals. Mutations are colored by their supporting evidence from WGS: only present as heterozygous in filia (grey), present in filia and subclonally in regenerants (red). **C)** Scatter plot of allele frequencies of subclonal mutations detected by WGS versus by amplicon sequencing of PCR products. Mutations are colored as in B). **D)** Allele frequencies of subclonal mutations in regenerants and parental animals, demonstrating that few mutations were already present in parental animals. **E)** Immunofluorescence for H3P marker of mitosis, on amputated tails equivalent in size to the fragments from which regenerant animals formed. **F)** Quantification of the fragment size and number of H3P positive cells in each fragment. **G-H)** Possible ways of organization of stem cell clones in the planarian body, symbolized by colored markers. **G)** Different clones are spread uniformly across the body. **H)** Clones are spatially separated. Pie charts show the distribution of colored clones in the cut tail and the rest of the body.

*De novo* mutations are expected to appear only on one chromosome; therefore, their allele frequency must be multiplied by the ploidy level to estimate the proportion of cells carrying the mutation. The median regenerant allele frequency of PCR-tested clonal F_R_ mutations was 1%, and the maximum was near 6% by both PCR and WGS, thus typically 3%, but as many as 18% of the cells of the respective regenerant carried the mutations detected subclonally in their germline and clonally in their offspring. This raises the question whether the mutations in the regenerants were already present at the time of the tail amputation. Notably, WGS detected supporting reads in the corresponding parental P_R_ samples for 6 of the subclonal mutations in the regenerant group, and by re-testing 32 F_R_ mutations in the P_R_ samples by PCR we found another two such SNVs. Altogether 8/46 regenerant mutations were detected in P_R_ animals, and these included those with the highest AF in the regenerants (Fig. 6D). This suggested that parental animals, even before regeneration, contained a diverse set of stem cells, and some of the detected *de novo* mutations in the filial generation resulted from the expansion of mutation-containing stem cell clones contributing also to germ cells in the regenerant animals. The remaining subclonal mutations in regenerants arose during regeneration. Altogether, the results of the regeneration-associated mutational analysis suggested that only a limitted number of stem cells was required to regenerate animals, thus we expected to see a low number of dividing stem cells in an amputated piece of tail. On the contrary, an average of ∼250 dividing stem cells was detected immediately after amputation in tail fragments, as determined by immunofluorescence of phosphorylated H3 histone (Fig 6E-F). H3P staining underestimates the actual number of stem cells in the fragment, as it labels only those cells that are dividing in a given moment. If we consider that the actual number of stem cells is even higher, we can conclude that certain stem cells contribute disproportionately to the regenerated body of the animal, including the germline.

## DISCUSSION

We present here the first draft genome assembly of *S. polychroa*. Using a strategy that integrates long-read PacBio and Illumina sequencing methods, we provided a relatively contiguous genome assembly on 8253 scaffolds. Using multiple theoretical methods and an experimental one based on flow cytometry, we estimated that the genome size of *S. polychroa* is around 588 Mbp. Allele frequency distribution of high-confidence germline mutations showed that all animals included in this study were triploid, indicating that no occasional sexual reproduction occurred in the analysed population of animals. Repetitive elements of *S. polychora* were divergent from those in the human genome, but similar to that of the related planarian species *S. mediterranea* [22]: we found a higher percentage of DNA transposons and unclassified DNA elements. DNA transposons tend to integrate in the proximity of coding genes, and can also be efficiently excised from the sites of integration, having an effect on the organization and expression of nearby loci [41]. Increased proportions of transposable elements might give rise to new allelic forms, which could be an important mechanism of diversification in a species that reproduces primarily by parthenogenesis. It has been shown recently that transposable elements can increase the levels of heterozygosity in inbred populations [42].

We also provide a transcriptome of *S. polychroa.* While 71% of transcripts mapped uniquely on the genome assembly, a fraction of transcripts mapped twice or multiple times on the genome, mostly due to repetitive elements, and some transcripts appeared to be chimaeras. Chimeric mappings can be signs of assembly fragmentation, but it is also notable that a relatively large fraction of chimeric transcripts marked spliced leader trans-splicing. This particular form of trans-splicing has been observed in a range of eukaryotes, including *Platyhelminthes* species [43] [32], and *S. polychroa* could potentially be a useful system to study this process.

Despite the fragmented nature of the bm_Spol_g1 genome assembly, it will enrich existing genomic information in the genus of planaria, and potentially provide opportunities for comparative genomic studies with related species.

In the mutational part of our study, we estimated the *de novo* mutation rate of *S. polychroa* and described its associated mutational profile. Described *de novo* mutations do not necessarily have a uniform origin. Presumably, some mutations can arise in somatic stem cells and persist at the subclonal level prior to the germline specification, while others form in differentiated germline, during gametogenesis. Both type of these mutations eventually become a parental germline mutation and a clonal mutation in the offspring. For this reason, the reported *de novo* mutations represent the mutational load of one generation, and allow for the estimation of the *de novo* mutation rate. The profile of *de novo* mutations resembled the “clock-like” SBS5 (single base substitution 5) signature in the COSMIC (Catalogue of Somatic Mutations in Cancer) database [44]. This signature correlates with aging and is considered a background mutagenic component both in human cells [45] and various other Eukaryotes [46]. We also described linage-specific germline mutations, common to all animals in this study. It would be interesting to understand the origin of these mutations and whether they are related to parthenogenesis as a mode of reproduction.

When animals passed through a single round of externally induced regeneration, the number of mutations increased three-fold and we detected an alteration of the mutational profile. This is the first estimation of genomic alterations related to the process of regeneration in any living organism. The caveat of this observation is that it was made on a limited number of sequenced samples and future investigations on a larger pool of samples and in different systems should provide support for the reported observation. The genome-wide analysis of mutation allele frequencies helped understanding the regenerative development of *S. polychroa*, with three notable findings. First, all of the clonal *de novo* mutations in *filia* were also detected in the regenerants, often with high (in cases, over 10%) contribution to the regenerant body. Most of these mutations likely arose during regeneration as they were not detected in the parent, and their spectrum differed from somatic mutations in control animals. Single stem cells therefore appear to have contributed to large fractions of the DNA content and cell number of the regenerated and fully grown animals. The ∼1% median regenerant allele frequency of clonal filial mutations implies a ∼3% contribution by each mutation-acquiring stem cell to the full body. Though the number of dividing stem cells in the amputated tail fragment was large, as indicated by the mitosis marker H3P, only a limited number of stem cells contributed to rebuilding the rest of the animal. This suggests that intraindividual selective pressures act on single cells, an evolutionary concept that has been described previously [47].

The second surprising finding was that certain clonal mutations in *filia* were present at similar, relatively high proportions in the cells of the regenerants and very few of them in their parent. Small number of such mutations, detected in the parental animal, suggested that the mutation-carrying somatic lineages made similar contributions to the cut tail and the remaining body of the parent, implying that planaria develop in a mixed mosaic manner, with cell lineages created by early divisions and marked by early somatic mutations contributing in similar proportions to different parts of the body (Fig. 6G). It is also possible that the similar contributions of mutation-carrying stem cells to the body and the tail resulted from the proportional cutting of spatially separated somatic clones (Fig. 6H), but we consider this less likely due to the similar contributions of several mutations to P_R_ and R animals. Though technically challenging, whole genome sequencing from several separate body segments, could decisively differentiate these possibilities. Somatic mosaicism has been described previously in planarians, analysing two genetic loci in animals with different reproductive strategies in the species *Dugesia subtentaculata* [48].

Finally, the observed mutation spectra were also revealing. The altered mutation spectrum upon regeneration confirmed that regeneration comes at a cost at the genome level. Importantly, this change in the spectrum was unique for regeneration, as lineage-specific germline mutations, shared by all animals, were more similar to the spectrum of the control *de novo* mutations, in accordance with the fact that *S. polychroa* does not reproduce by fission. We can only speculate as to the cause of the abundance of C>T and T>C mutations in the regenerated *filia*. The same mutation classes dominate the SBS spectra of mismatch repair deficient cancer cells alongside oxidative damage-induced C>A mutations, and the C>T and T>C transition mutations are thought to primarily result from polymerase errors [49]. It is thus possible that fast cell proliferation during planarian regeneration is accompanied by a reduced efficiency of DNA mismatch repair, resulting in the altered mutation rate and spectrum.

Our results highlight certain parallels between planarian regeneration and mammalian biology. The similar contribution of cell lineages to the body and the tail of planarian can result from early embryonic cell mixing also evidenced by patterns of chimeric mammals, or by extensive cell migration also typical of mammalian development [50]. Tissue regeneration is best seen in the liver of mammals, and the regrowth of relapsed tumours following surgery can also be considered an example of a regenerative process. Genome alterations accompanying these processes are relevant to subsequent tumorigenesis or the development of resistance, and the planarian system can be a useful model for understanding the costs of tissue repair at the genome level.

## MATERIALS AND METHODS

### Animal husbandry

*S. polychroa* was a kind gift of the Aboobaker lab and the asexual biotype of *S. mediterranea* was a kind gift of the Rink lab. Planarians were grown in 0.05% Instant Ocean at 20°C and fed organic calf liver once per week. Animals were starved two weeks at least before genomic DNA extractions or one week at least before other experiments.

### Genomic DNA extraction and sequencing

For long-read PacBio sequencing we used the previously published protocol for planarian genomic DNA extractions [22]. For Illumina sequencing of single animals, we introduced a slight modification in the extraction protocol. Briefly, animals were treated with 0.5% N-acetyl-L-cysteine (NAC) pH 7.5 in agitation to strip off the mucus. NAC was replaced with a lysis buffer: 4 M guanidinium thiocyanate, 25 mM sodium citrate, 0.5% (w/v) N-lauroylsarcosine, 7 % v/v β-mercaptoethanol and animals were swiftly disrupted with a motorized pestle for several seconds. This allowed for higher yield of genomic DNA without shearing. Animals of average size: 1.5-1.8 cm, were lysed in 1 ml of lysis buffer. The protocol further follows that of Grohme *et al*, including the post-purification step of mucopolysaccharide removal with cetyltrimethylammonium bromide [22]. Purified DNA was resuspended in 1x TE and DNA quality was assessed by NanoVue, Qubit and 0.7% agarose gel electrophoresis. Only samples with NanoVue readings 260/280 ∼ 1.8; 260/230 ∼ 2, minimal concentration (estimated by Qubit) of 25 ng/μl and no indication of DNA degradation on an agarose gel were sequenced. Library preparation and sequencing of Illumina datasets (2×150 bp, paired end) were performed by Novogene (Beijing, China) using Illumina NextSeq 2000 instruments, obtaining approximately 50X genome coverage. Long read library preparation and sequencing was performed on PacBio Sequel machines by Novogene, resulting in 18.04 Gbp read length on average, with 20x genome coverage.

### Genome size estimation by FACS

In total 5 samples containing 6 animals each were collected for genome size estimation. *Schmidtea mediterranea* was used as a reference sample of known genome size. Frozen samples were homogenized briefly in Galbraith buffer (45 mM MgCl2, 30 mM Sodium citrate, 20 mM 3-(*N*-morpholino) propanesulphonic acid (MOPS), 0.1% v/v Triton 100x) with a motorized pestle. Homogenized samples were passed sequentially through a 70 μm and then 40 μm mesh. Nuclei were pelleted by centrifugation, resuspended in 1X PBS with 50 μg/ml propidium iodide, 200 μg RNAse A, and incubated for 2 hours on ice, prior to data acquisition. Flow cytometry data were acquired on BD FACSVantage SE (Becton Dickinson) and analyzed using FlowJo software Macintosh version 8.1.1 (Tree Star).

### RNA extraction and sequencing

Total RNA was extracted as described previously [51], with a modified post-purification step. Namely, after Trizol extraction and DNAse treatment, RNA was ethanol precipitated to avoid loss of short transcripts. 4 animals of different size were used for extraction of total RNA. RNA quality was estimated by NanoVue. Strand-specific library preparation and sequencing (100 bp, paired-end) was done by BGI (Hong Kong, China) using DNBSeq.

### Immunofluorescence

Tails of animals were amputated and inactivated immediately in 2% HCl for 5 minutes. Three hours fixation in Carnoy fixative (60% ethanol, 30% chloroform, 10% acetic acid) followed. Carnoy was replaced with methanol and after one hour, samples were rehydrated and blocked in a blocking solution (1 x PBS, 0.3% Triton X-100, 5% horse serum), followed by incubation with the primary antibody (anti-H3P, Cell Signaling, 1:200) overnight. After overday washes in PBST (1 x PBS, 0.3% Triton-X100), samples were incubated with the secondary antibody (Alexa 488, Jackson’s lab, 1:400) overnight. Samples were mounted on a glass slide in Mowiol.

### *De novo* genome assembly

The draft genome was constructed from PacBio long reads using Canu v1.8 [52] with parameters optimized for relatively low coverage (correctedErrorRate = 0.105, corMinCoverage = 0, corMhapSensitivity = “high”, corOutCoverage = 10000, corMaxEvidenceErate = 0.15). Three rounds of polishing were performed by Arrow 2.3.3 (https://github.com/PacificBiosciences/pbbioconda) followed by one polishing round with Pilon 1.23 [53], both with default parameters. The Illumina dataset used for Pilon polishing was error corrected by Quake 0.3 [54] with a k-mer value of 19. Redundant contigs were removed using a multi-step purging strategy. First we ran the purge_dups pipeline [55], utilizing Illumina reads aligned with minimap2 [56] and using coverage cutoff values of 5, 40 and 70. Next, further haplotig reduction and first-level scaffolding using Illumina and PacBio reads was achieved by running all three steps of redundans.py [57]. The remaining haplotigs were removed using a custom script. Briefly, first we performed an all-to-all mapping using minimap2 with parameters -x asm5 -D -n 18 –secondary=no and filtered the resulting alignments that were not one-to-one mappings, had a secondary chain score larger than half of the primary chain score, had more than 60% divergence in the mapped region, had mapping qualities less than 60 or that were overlapping with one single repeat element on more than 40% of the aligned length. Next, we removed the filtered hits (which are present on two different scaffolds) from the shorter containing scaffold. Final scaffolding was done using the PCFL *de novo* transcriptome assembly with L_RNA_Scaffolder [58]. As the last step, we removed all scaffolds shorter than 1 kbp, having a GC ratio over 40% or an average Illumina coverage less than 10. All scaffolding rounds were followed by gap closing with GapCloser 1.2 [59], resulting in the final draft genome referred to as bm_Spol_g1.

### Transcriptome assembly

RNA-sequencing reads were assembled with Trinity [60] using default parameters. The raw *de novo* transcriptome assembly was annotated by the Trinotate [61] pipeline. The probable protein coding fraction of the transcriptome assembly was generated by filtering the raw transcripts for the presence of BLASTX hits against the Uniprot/Swissprot database, or the presence of at least one Pfam domain, or a minimal length of 100 bp. Gene annotations were determined using the raw transcriptome assembly by GMAP [30]. For some analyses, we further filtered the transcriptome dataset to obtain the longest isoforms of each gene (PCFL set).

### Genome characterization

Genome size and completeness was estimated using four methods: k-mer based estimations were obtained using GenomeScope2 [62]; the coverage-based calculation used the Lander-Waterman equation (G = LN/C, where G is genome size, L is read length, N is the number of reads and C is the genome-wise average coverage); we also utilized BUSCO [25] with the eukaryota_odb10 and the metazoa_odb10 datasets. Cellular ploidy was determined by observing the allele frequencies of germline mutations present in all analysed animals. Repetitive element families were predicted by RepeatModeler 2.0.1, and the resulting repeat families were detected in the draft assembly by RepeatMasker 4.0.9 [63]. Genes were annotated by remapping the transcripts in the PCFL set onto the final genome assembly by GMAP [30]. Multi-mapper, duplicate and chimeric transcripts were manually curated by their overlaps with repeat elements, by the presence of spliced-leader trans-splicing leader sequences as determines by the SLIDR/SLOPPR pipeline [31], and the lengths and relative positions of the multiple mapping sites. The scaffold containing the mitochondrial genome was manually curated, and mitochondrial genes were annotated with the MITOS webserver [64]. Synteny blocks between *S. polychroa* and *S. mediterranea* were determined by mapping the PCFL transcript sets of both species onto their genome assemblies (bm_Spol_g1 and dd_Smes_g4, respectively) by GMAP, and extracting homology information from the PlanMine API [29]. To ensure that only clear homologies were used, we only selected orthologous pairs with one-to-one homology, and only considered those *S. polychroa* transcripts that were masked on less than 10% of their length, had a mapped length of more than 300 bp, and were not inside another gene. The dd_Smes_g4 scaffolds were mapped using minimap2 onto the chromosome-level assembly of *S. mediterranea* [33].

### Mutation calling

Illumina sequencing datasets from the parental, regenerant and *filia* animals were aligned against the bm_Spol_g1 draft reference genome with BWA mem [65]. Lineage-specific mutations (i.e., pre-existing events shared between a parent animal and its progenies) were found by the HaplotyeCaller tool from the GATK 3.8 package [35]. *De novo* mutations in *filia* were found with IsoMut [11] and HaplotypeCaller. HaplotypeCaller hits were postfiltered in each case by extracting the number of supporting reads in each potential heterozygous site using the pysam Python library, and only those events were retained that had more than 7 supporting reads in the candidate samples, but no more than 1 read in any other, having a mean coverage of 25-130x in the assessed samples, and had an average mapping quality over 40. IsoMut was initially ran with permissive settings (min_sample_freq = 0.15, min_other_ref_freq = 0.9, cov_limit = 20), and potential hits were filtered to have at least 7 supporting reads and coverage between 25 and 130x. All candidate mutations were manually verified in the Integrative Genome Browser 2.12 [66].

### Validation of mutations

Subclonal mutations detected in regenerant animals were validated by PCR. Only those candidate events were considered that were positioned in uniquely mappable +/-250 bps regions as assessed by BLASTn. Primer pairs were designed to amplify regions surrounding each of the unique mappable events (Table S8) by PCR, and the resulting amplicons were mixed and sequenced on an Illumina MiSeq instrument (Cogentech, Milan, Italy). The resulting 2×150 bp paired-end reads were trimmed with cutadapt [67] to remove adapters, merged with FLASH2 [68] using parameters -M 150 and -x 0.05, and aligned against the bm_Spol_g1 assembly with BWA mem. Allele frequencies of mutations were determined manually by inspecting the mutation sites in the Integrative Genome Browser.

#### Data and code availability

The assemblies and the aligned Illumina datasets presented in the study were deposited in the European Nucleotide Archive (ENA), under the accession ID PREBJ58888. The workflows used in the analysis, also including any custom scripts, are accessible on GitHub: https://github.com/szutsgroup/Schmidtea_polychroa_regeneration.

## ACKNOWLEDGEMENTS

We thank Prof. Marco Foiani for continuous support of this research project. This work was funded by the generous donation of the Suma-Nesi family.

## AUTHOR CONTRIBUTIONS

J.V. conducted experiments. A.P. performed all bioinformatic and quantitative analyses. A.P., J.V. and D.S. discussed, wrote and edited the manuscript. J.V. supervised the project.

---

**Figure S1.**
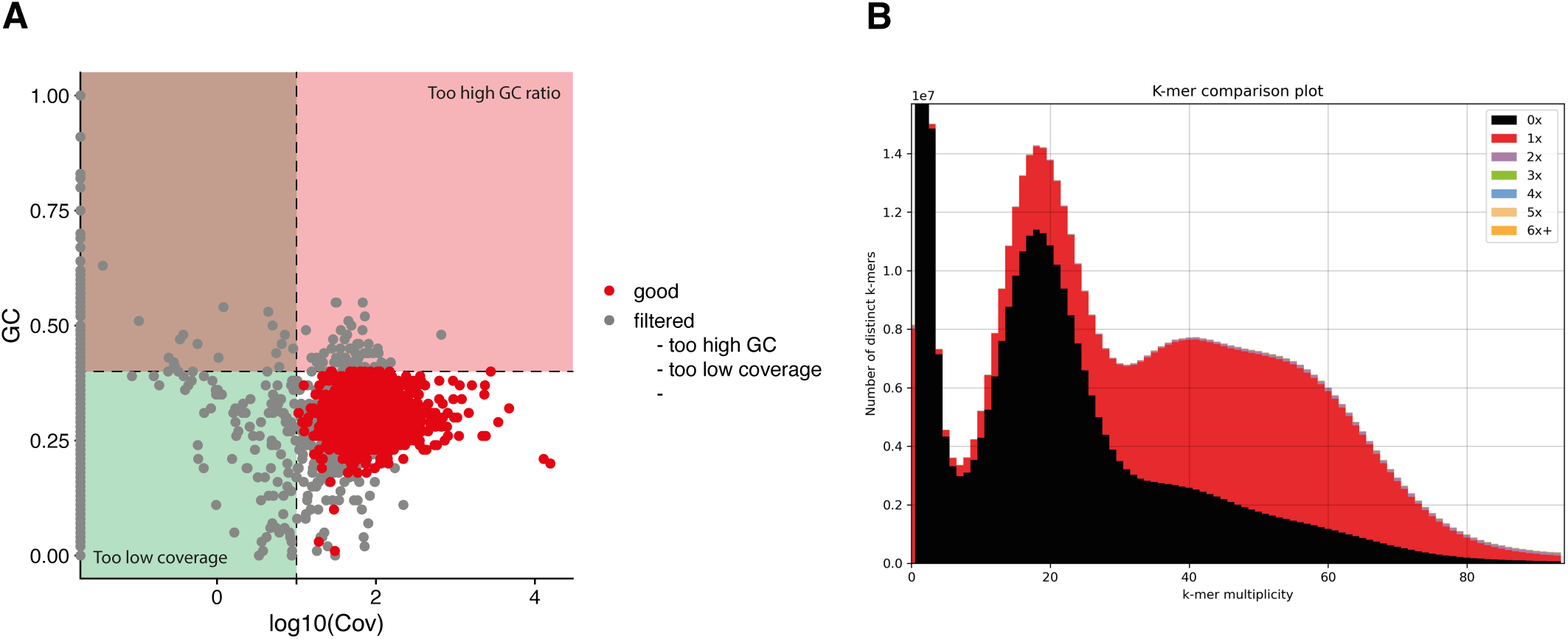
Genome assembly quality control. **A)** Scatter plot of GC ratios (Y axis) and Illumina coverages (X axis) of purged scaffolds. Grey points refer to scaffolds that were filtered either because they had GC ratios over 40% (pink region) or because they had coverages less than 10x (green region). Only contigs marked by the red markers were included in the final bm_Spol_g1 assembly. **B)** KAT profile of the final assembly. The three peaks suggest that the S. polychroa genome is triploid, the relatively large black region belongs to the genomic regions missing from the assembly, and the virtual lack of k-mers present more than once suggest that the assembly is appropriately purged.

**Figure S2.**
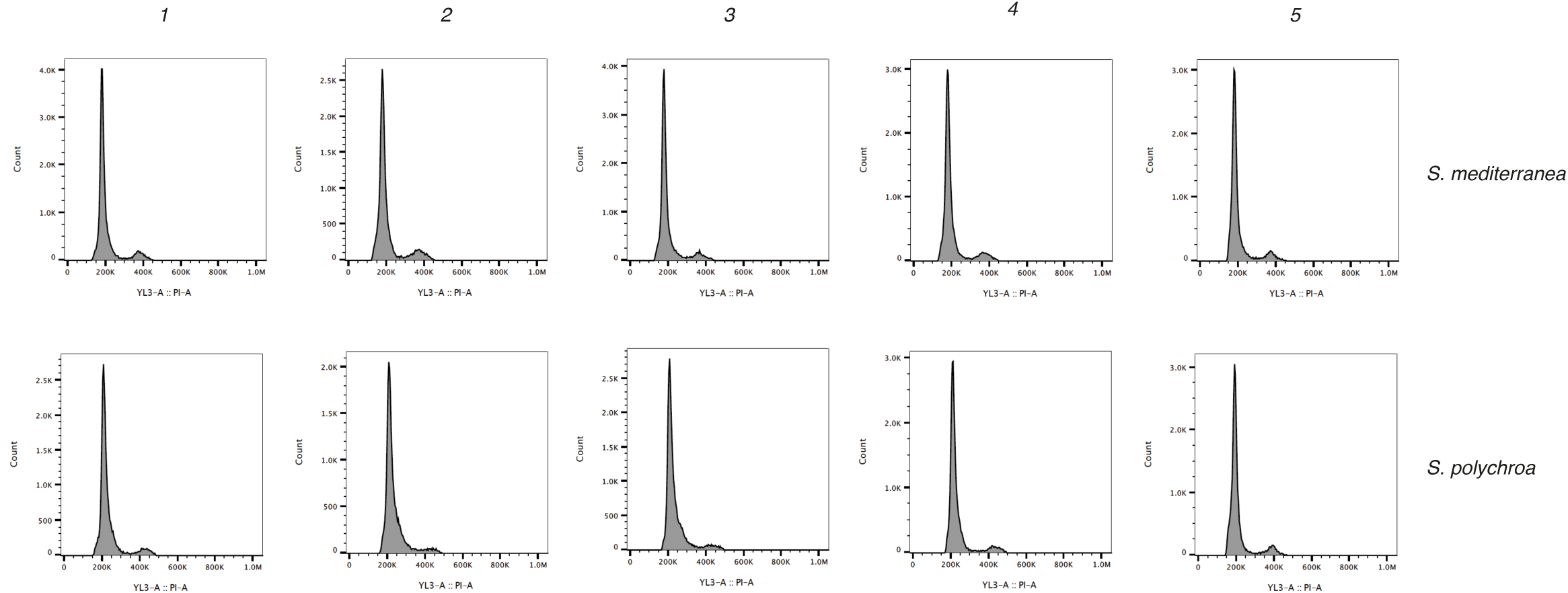
FACS profiles of all 5-5 *S. polychroa* and *S. mediterranea* samples for the genome size determination experiment.

**Figure S3.**
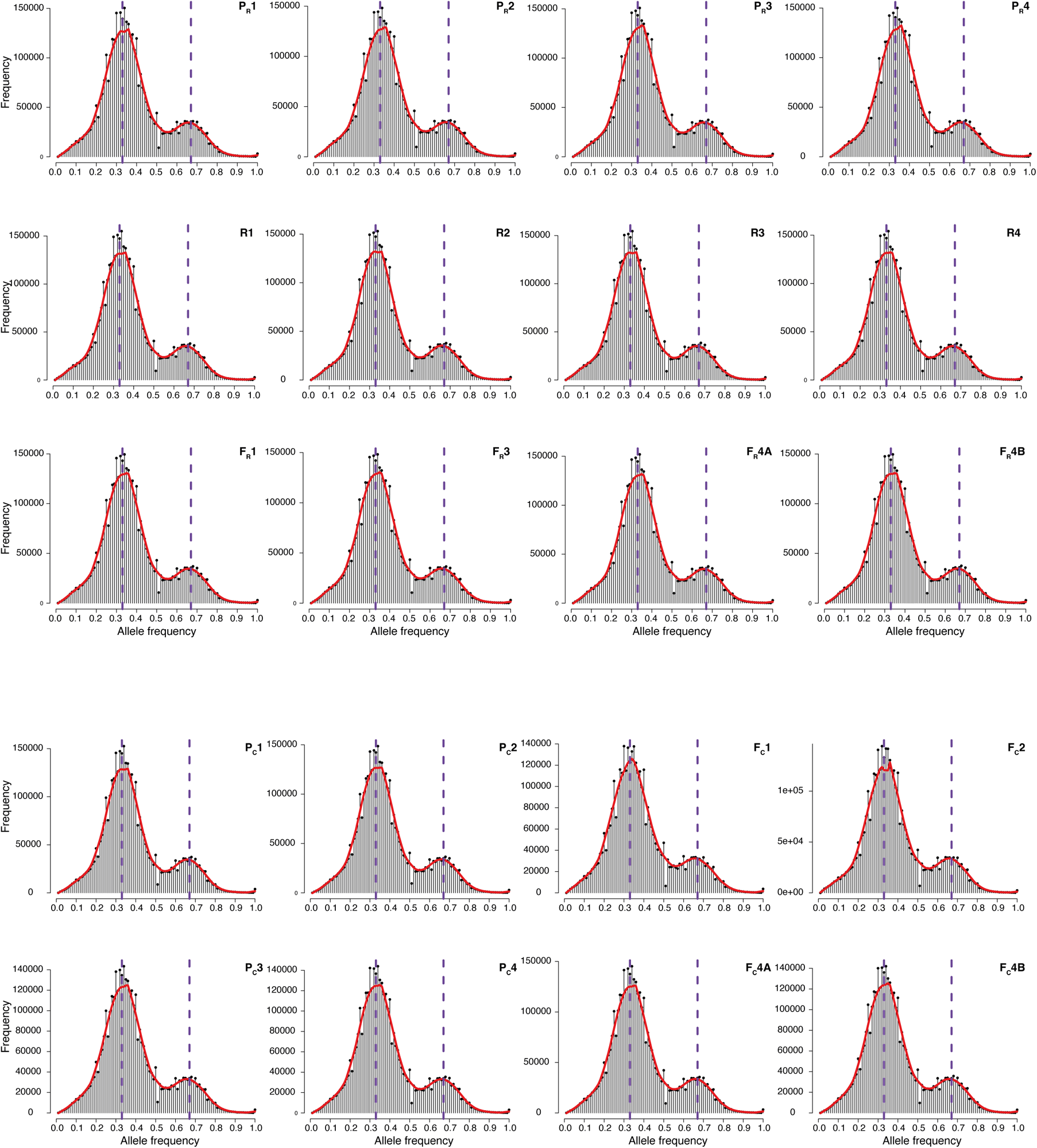
Annotated map of the mitochondrial genome of *S. polychroa*. Blue arrows: protein coding genes; Purple arrows: ribosomal RNA genes; red markers: tRNA genes.

**Figure S4.**
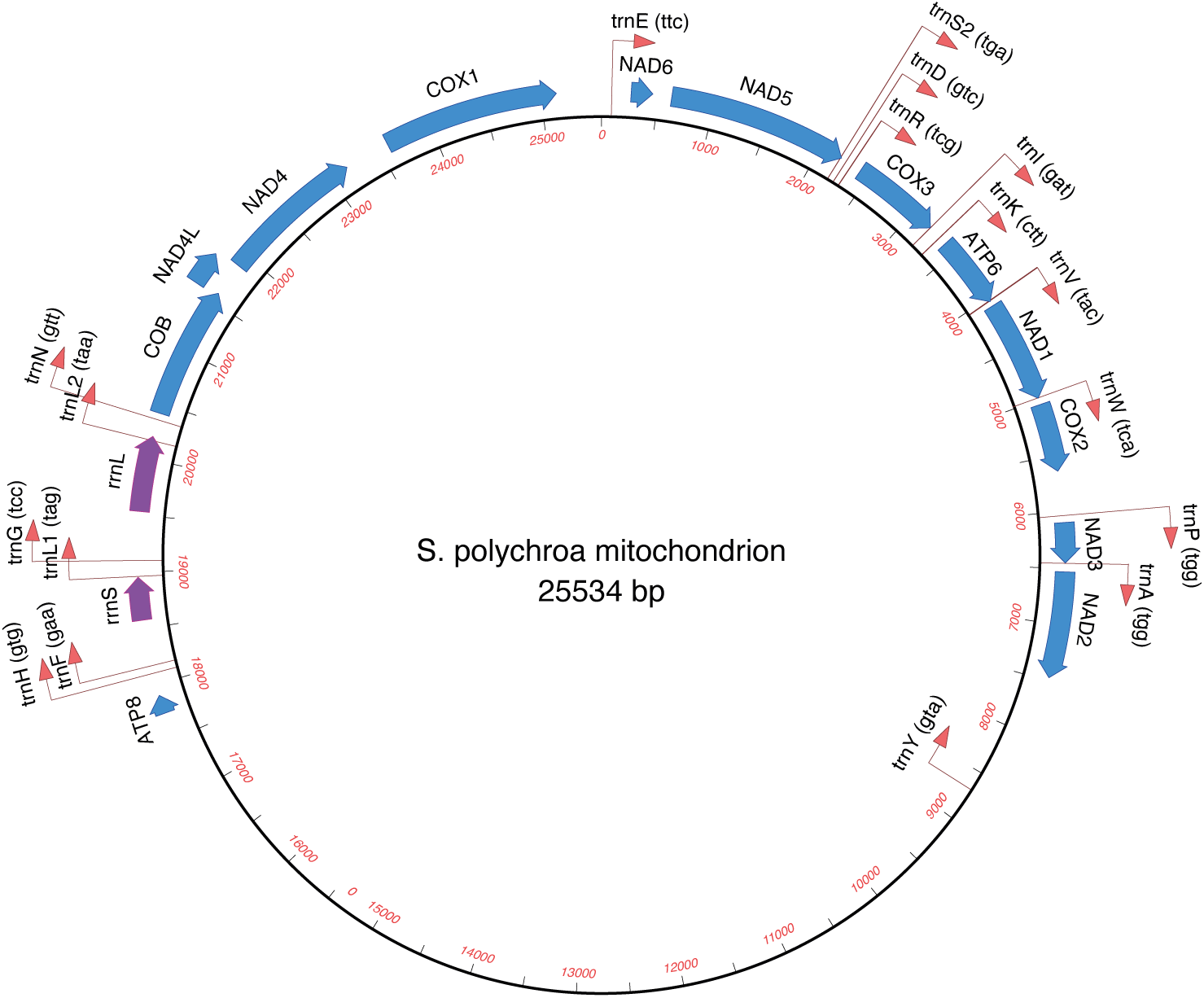
**A)** Bowtie2 alignment statistics after mapping the RNA-seq reads used for the assembly onto the bm_Spol_tr1transcriptome assembly. **B)** Histogram of mean coverages of the RNA-seq back-mapping. **C)** BUSCO outputs of the raw and the probable protein-coding fraction (PCF) filtered bm_Spol_tr1transcriptomes, compared to the dd_Spol_v4_pcf transcriptome downloaded from PlanMine. **D)** Venn diagram of transcripts affected by the three filtering factors for the PCF set generation.

**Figure S5.**
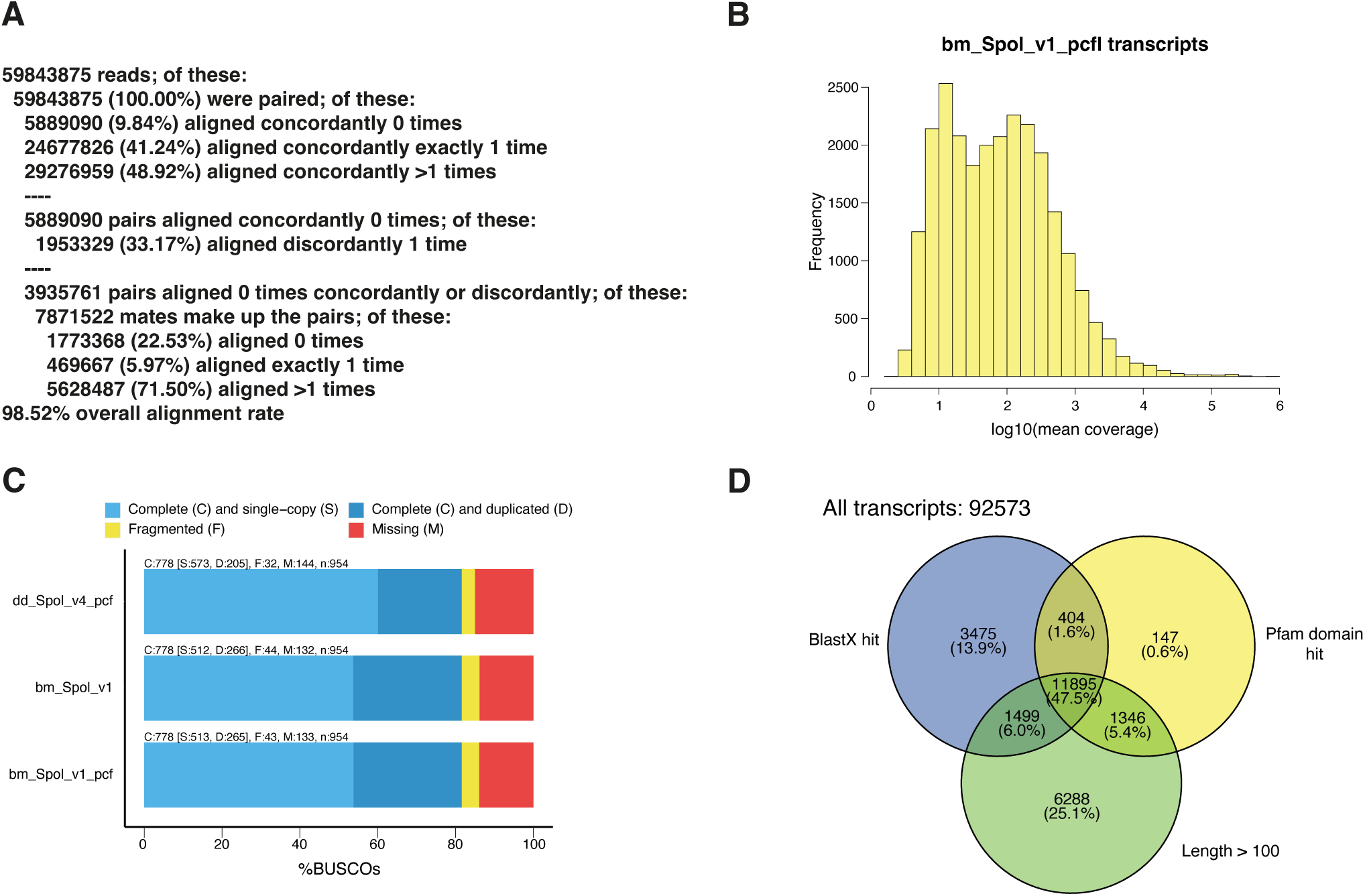
Spliced-leader sequence in *S. polychroa***. A)** Alignment of the spliced-leader RNA genes found in *S. polychroa* and the SL1 gene sequence from *S. mediterranea*. The colored segments mark stem regions. **B)** Secondary structure of the *S. polychroa* spliced-leader RNA. The colored segments are identical to the regions on A). The splice-site is marked wirt a black arrow.

**Figure S6.**
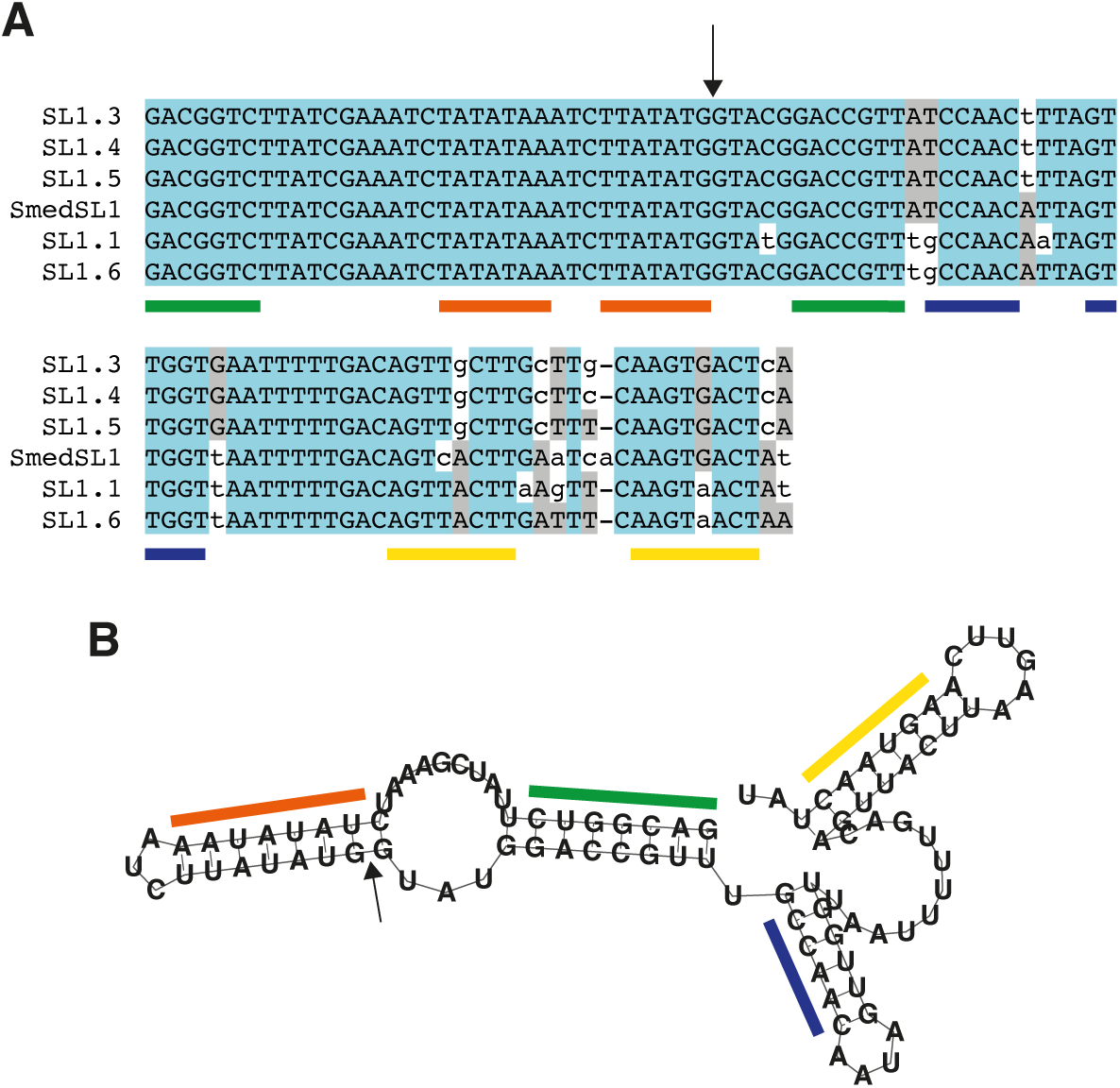
**A)** Counts of *S. mediterranea* scaffolds corresponding to each *S. polychroa* scaffold, determined using one-to-one homologous gene pairs from PlanMine. **B)** Distribution of synteny block lengths, defined as the start of the first and the end of the last gene in a block. **C)** Detailed analysis of regions corresponding to observed rearrangements between the two *Schmidtea* species. Only breakpoints with at least two genes on both sides in the *S. polychroa* assembly were considered, and misassemblies were defined as breakpoints with zero coverage in both Illumina and PacBio alignments.

**Figure S7.**
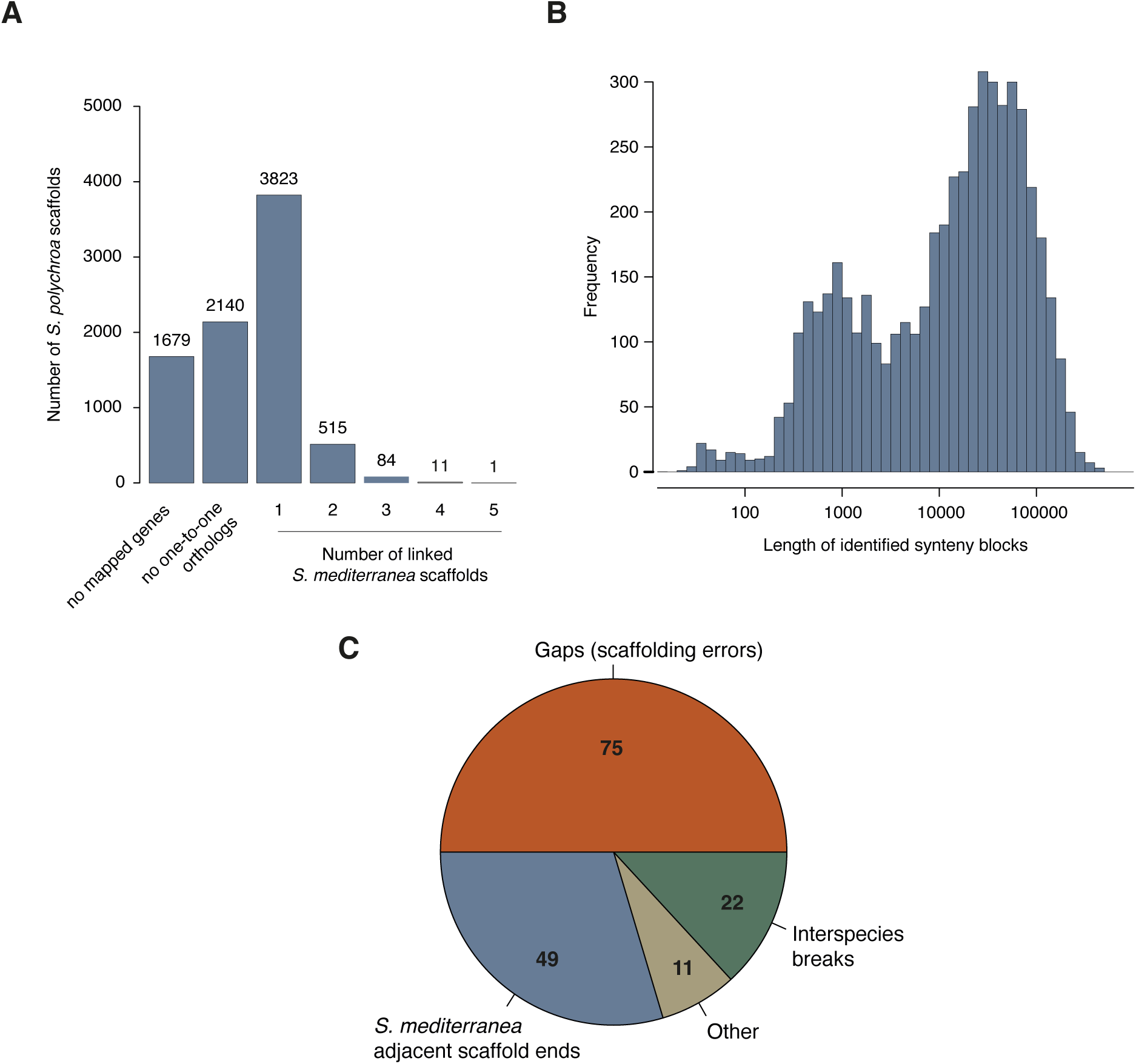
Substitution spectra of *de novo* mutations in the control and regenerated groups, and of germline mutations present in all animals.

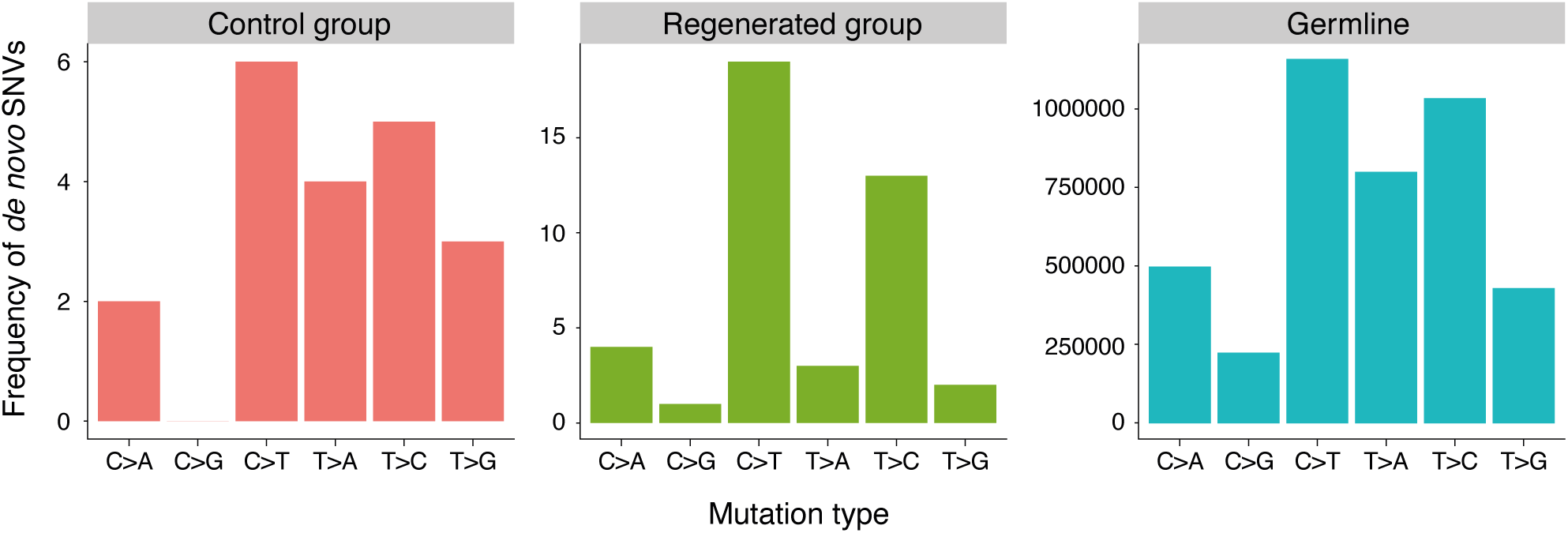

